# Functional MRI Investigation on Paradigmatic and Syntagmatic Lexical Semantic Processing

**DOI:** 10.1101/2020.01.11.899831

**Authors:** Songsheng Ying, Sabine Ploux

## Abstract

The word embeddings related to paradigmatic and syntagmatic axes are applied in an fMRI encoding experiment to explore human brain’s activity pattern during story listening. This study proposes the construction of paradigmatic and syntagmatic semantic embeddings respectively by transforming WordNet-alike knowledge bases and subtracting paradigmatic information from a statistical word embedding. It evaluates the semantic embeddings by leveraging word-pair proximity ranking tasks and contrasts voxel encoding models trained with the two types of semantic features to reveal the brain’s spatial pattern for semantic processing. Results indicate that in listening comprehension, paradigmatic and syntagmatic semantic operations both recruit inferior (ITG) and middle temporal gyri (MTG), angular gyrus, superior parietal lobule (SPL), inferior frontal gyrus. A non-continuous voxel line is found in MTG with a predominance of paradigmatic processing. The ITG, middle occipital gyrus and the surrounding primary and associative visual areas are more engaged by syntagmatic processing. The comparison of two semantic axes’ brain map does not suggest a neuroanatomical segregation for paradigmatic and syntagmatic processing. The complex yet regular contrast pattern starting from temporal pole, along MTG to SPL necessitates further investigation.

## I. INTRODUCTION

Some functional magnetic resonance imaging (fMRI) investigations on word-meaning processing in the human brain have correlated semantic embedding vectors with neural activity recording. Especially it probes into the representational and/or operational functionalities of neural regions. Previous studies [1]–[6] revealed a distributed pattern of cortical areas’ correlation with word semantic models. The relatively well-modeled voxels are typically located in the temporal, parietal and frontal lobes. Studies show that context length [5], semantic category [2], [7], conceptual concreteness [8] and perceptual properties [9], [10] are all related factors to differentially elicit semantic activities in different lobes. Yet, despite abundant literatures on theoretical semantic network architecture[11], no concluding evidence enlightens each cortical region’s functional role in the processing.

Classical view [12] supported a temporal construction of semantic processing. To bridge the temporal-centric semantic processing theories and the found distributivity, a structure of *semantic hub* [13], [14], located in the bilateral anterior temporal lobe (ATL), acting as a centralized location that hosts and/or processes semantics, is by far consistent with human neural modeling studies [15], anatomical analysis [16] and pathological observations [17]. For example, in patients with semantic dementia (SD), the ATL atrophy and hypo-metabolism lead to a conceptual deficit. These patients develop a specific loss of capability to use distinctive features (such as stripes for zebras) necessary to identify and characterize a concept. So that, they will fail to name specific categories (eg. zebra) and would use similar or hypernymous categories (eg. horse) instead. The semantic hub’s involvement in linguistic and non-linguistic semantic tasks indicates a generalized supra-modal or amodal concept network storage in ATL, beyond word meanings. It is said to accept inputs from a specific semantic neural component and activates reciprocally other components to evoke a complete mental representation of a word. Therefore, the hub must support a holistic structure of all words with at least minimal information. [18] suggests that the left ATL is a non-syntactic conceptual hub involved in semantic composition. The convergence of ATL’s relatedness to semantic composition of conceptual attributes, to the preservation of hypernymous or similarity links (zebra–horse) and to the loss of the distinctive features in case of lesion to this region, needs to be further explored. As for the other components in the semantic processing system, they are either responsible for a modality specific semantic information stream or a semantic control/computation function.

An analogy in structural linguistics for this two-fold semantic processing is the paradigmatic and syntagmatic axes proposed by De Saussure, Jakobson and Halle. The two-axis proposition is best illustrated by an example in Table I. To render the paradigmatic *absentia* candidates, the responsible neural structure should hold at least a local lexical similarity comparison in the paradigmatic sense, and sort out the most similar words. The syntagmatic axis looks for semantic associations mainly by collocation. The functional association of syntagmatic semantics with the neural components found active in the aforementioned previous studies needs to be further explored. In addition, Saussure’s theoretical proposal is completed by a psycholinguistic meta-analysis by [20] on two dissociated groups of aphasics each associated with one semantic axis, namely selection-deficits for paradigmatically impaired aphasics, and contexture-deficits for syntagmatically impaired ones, implying potentially different cortical loci for the processing of two axes.

**TABLE I.**
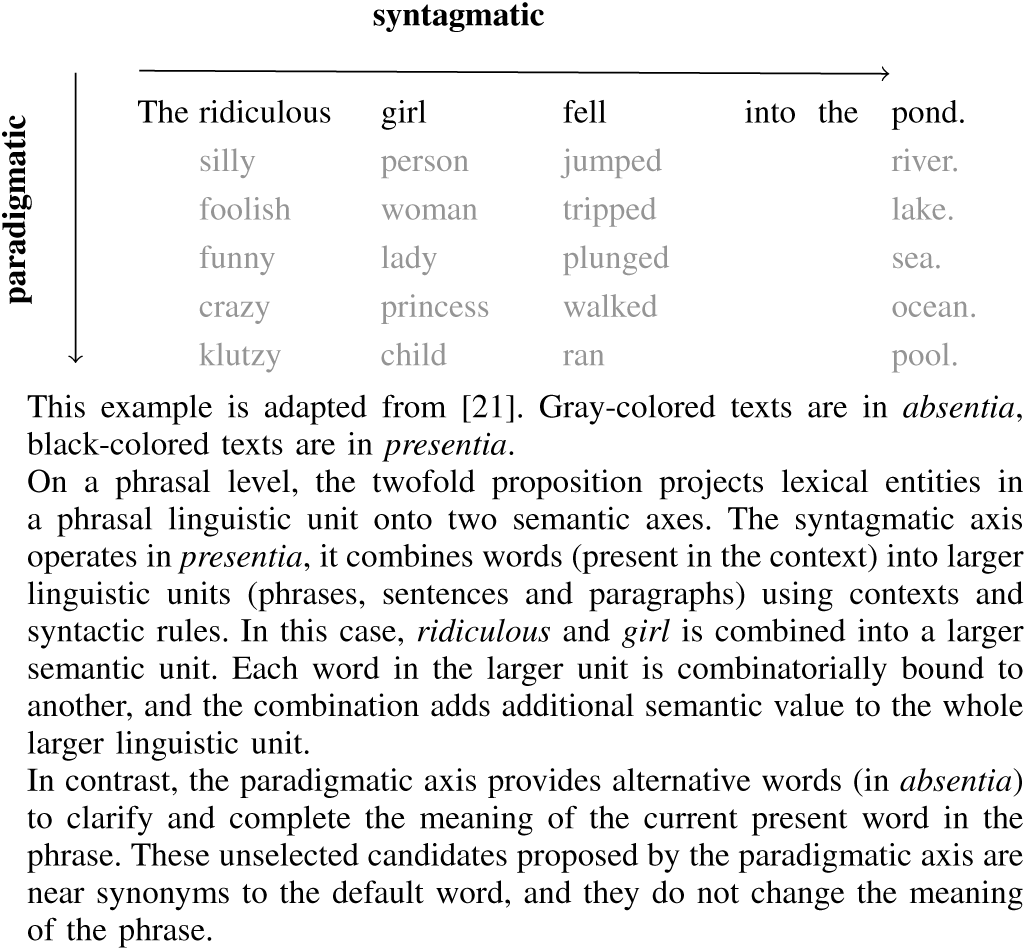
An Example of Syntagmatic and Paradigmatic Axes

This study leverages the contrast of paradigmatism and syntagmatism to investigate the nature of each cortical region’s involvement in lexical semantic processing.

## II. METHOD

This study hypothesizes on a representational and/or operational account that the paradigmatic and syntagmatic axes engage different yet not necessarily dissociated neural structures during semantic processing. Consistent with other works using fMRI encoding experiments [1], we propose an investigation by modeling neural activity with semantic embedding features, separately for the paradigmatic and syntagmatic axis, to potentially segregate the respective responsible neural structures.

The experiment is conducted with priorly collected data from the framework project “Neural Computational Models of Natural Language” (Principle Investigators: John Hale and Christophe Pallier). The stimuli materials preparation, participant recruitment, fMRI experiment and fMRI data preprocessing are carried out in [22].

Following the standard fMRI encoding procedure, the first step is to model the time-sequences of the administered stimuli received by experiment participants during the fMRI sessions. The external stimuli model will rely on paradigmatic and syntagmatic representation models (semantic embeddings), which themselves are abstract vectorial representations of semantic values in a high-dimensional space. The second step consists of voxel-level regression analyses which map the theoretical stimuli temporal signals onto actual fMRI measured brain activities. The brain activities are modeled as a temporo-spatial 4-dimension data. In MRI recordings, the brain volume is divided into small 3D volume units, namely *voxels*. Each voxel has a time series of spatially aggregated brain-activity measurement within its volume. A voxel-wise regression model is a mathematical function mapping the temporal semantic sequences onto the brain activity signal space.

The result analysis relies on each voxel model’s predictive power provided with its training feature dataset. Stronger the model performance (or the model performance improvement compared to another model), the stronger the correlation between the neural activity in that voxel and the semantic and non-semantic feature contained in the training set.

### A. Participants

20 French native speakers (11 females, average age of 24.5 years-old, range 18–39 years-old, right handed according Edinburgh’s inventory [23] adapted for French, averaged score 0.903, range 0.375–1, without antecedent neurological or psychiatric disorders) were recruited from NeuroSpin’s volunteer inventory. The recruited participants declare to have not been exposed to any material related to *The Little Prince* within last 5 years, including written books, audiobooks, films in any languages, and to be unable to clearly recall the story of the book.

### B. Procedure

After being received by a researcher and informed with the experience protocol, the participants are first examined by the lab doctor, then invited to scan an anatomical MRI which lasts 8 minutes. In the scanner, the participants are able to see the instructions and visual aides displayed on the screen through a mirror. During the anatomical scan, two illustrations from the first two chapters of *the Little Prince* [24] are presented to prepare the participants. A sound test phase follows: the introductory paragraphs are administered to the participants under the noise of MRI scanner. The auditory stimuli is adapted from the audiobook [25]. Once the sound adjusted, the functional imaging sessions begin.

For the comfort of the participants and their concentration during listening comprehension, the audiobook is divided into 9 blocks, so that each block lasts at most 15 minutes. For each chapter, the reading of chapter title is removed from the audio, and 3 seconds of silence is added. Within each block, the participants listened passively with eyes closed, to introduce the minimum amount of perturbation to the BOLD signal. At the end of each block, they were tested with comprehension multi-choice questions displayed on the screen and responded orally. Additionally, they were required to answer “comprehension questions” necessitating finer and deeper reflexions or to retell the synopsis of the past block. 4 or 5 blocks are played in the morning, and the rest in the afternoon. A final block is added, where the audio stimuli is comprised of normal French sentences and acoustically deformed sentences to localize language-processing related areas.

### C. MRI Acquisition and Preprocessing

Structural and functional data are acquired on a whole-brain 3-Tesla Siemens scanner at NeuroSpin. Structural images are collected in 154 axial slices with 1 mm isotropic voxels. Functional images are collected with multi-echo EPI sequence (3.159^3^*mm*^3^, TR = 2000 ms). Multi-echo sequence is used to boost BOLD and non-BOLD component identification based on TE dependence, resulting in a higher signal-to-noise ratio in the final image rendering [26].

MRI data is analyzed with Multi-Echo Independent Component Analysis (ME-ICA) toolkit (https://github.com/ME-ICA/me-ica). Raw structural data and 4-D time-series data grouped by TE are supplied to render de-noised sMRI and fMRI, which are also spatially warped in Montreal Neurological Institute (MNI) template.

### D. Paradigmatic Embedding

The paradigmatic embedding is built based on a French version of WordNet [27], [28], WOLF [29], a knowledge graph of words and abstract concepts. WordNet models synonym groups (*synset*) as vertices in a graph, and various semantic relations as edges connecting two vertices. The explicit semantic relations in WordNet are hand-coded by linguists. WOLF is a machine translated version of WordNet, manually verified and corrected by its authors.

With WordNet-alike knowledge bases, it is possible to compare pure taxonomical semantic similarity. Therefore, the knowledge base supports a modeling implementation of the paradigmatic axis. Such speculation is backed by a meta-analysis [30]. [30] collected a set of proposed distance metrics on WordNet then compared the word-pair ‘proximities’ against human similarity/relatedness judgement data (word-pair proximity ranking task). The results show that with WordNet it is possible to derive paradigmatic-specific semantic information. [31] uses WordNet retrofitting to enhance distributional embeddings. With the same ranking task and benchmarks, the enhanced versions perform better in paradigmatic tests and significantly worse in syntagmatic settings.

In WordNet-like knowledge bases, some of the semantic relations are paradigmatic: synonymy, hypernymy, hyponymy, the relation where an adjective is a participle of a verb (*exhausting*–*exhaust*), adjective having similar meaning (*exhausting*–*effortful*) and adverb deriving from adjective (*essentially*–*essential*). Other relations are syntagmatic: collocations, meronymy/holonymy (*ceil*–*house*), entailment/causality (*sunset*–*milky-way*). To build the paradigmatic embedding, We use [32]’s algorithm to transform the tree-like knowledge base consisting of *synsets* into a lexical semantic embedding, using a sub-graph of WordNet with only paradigmatic edges.

Consider vertices, which are more closely and densely connected by paradigmatic edges, more proximate in the paradigmatic sense. The algorithm implements first an infinite random walk to compute the graph distance between every word-pair with a discount factor for longer paths. In Eq. 1, *M* is the adjacency matrix of the knowledge graph, *γ* is a decay parameter weighting how longer paths are dominated by shorter ones. *M*_*G*_ is the convergent limit of the iterative process, which is the final state of the random walk. Then a normalized positive point-wise mutual information transformation is applied to reduce noises induced by unbalanced word occurrence frequency. Finally a principle component analysis (PCA) is applied for dimensionality reduction to render the final embedding output. The number of PCs are determined in the later feature engineering stage.

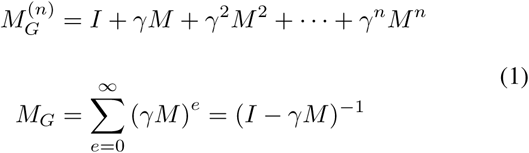

### E. Syntagmatic Embedding

The syntagmatic embedding is based on DepGloVe [33], a GloVe-alike [34] distributional semantic models [35]–[38], constructed based on the *distributional hypothesis* [39] that words having similar context have as well similar meanings. However, this consideration also brings close words consistently co-occurring in same contexts, such as *teacher* and *student*. In several analyses [30], [40], a set of pre-trained corpus-based distributional word-embeddings including GloVe are shown to always mix paradigmatic and syntagmatic axes into the actual embedding.

Since GloVe-like co-occurrence-based statistical distributed representation models contain both paradigmatic and syntagmatic information, it remains as an ideal source for syntagmatic information extraction as there exists no, to our knowledge, a pure syntagmatic embedding.

Under a linear-additive approximation of the mixture of the two semantic axes, a syntagmatic embedding can be extracted from a GloVe-like embedding by subtracting detectable paradigmatic counterpart. Suppose the abstract paradigmatic semantic relations modeled by the mixed embedding is also accounted by the paradigmatic embedding constructed in the previous section, then we can use the projection of the paradigmatic embedding onto the mixed space to reveal the paradigmatic component and separate the two semantic sub-spaces.

Consider `MIX`, `PAR` and `SYN` in Eq. (2) as 2D matrix representations respectively for the co-occurrence-based Dep-GloVe, the paradigmatic embedding built with WOLF, and the syntagmatic embedding to be constructed. The lexicons of the three embeddings are aligned such that each row with the same index from three matrices represents one same lexicon unit. *F* is a transformation matrix learned with general linear model, of which the computational objective is to minimize the L-2 norm of `SYN`.

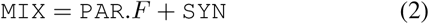

### F. Feature Engineering

To construct stimuli-based temporal sequence features, the audiobook and the original text of *the Little Prince* serve as the base. The onset and offset of each word in the audio document is aligned with the text. To aggregate the semantic value of various inflected French words, stemming is performed for source corpora and *the Little Prince* with `spaCy` and `FrenchLefffLemmatizer` [41]. The lemmatized text is manually verified.

Five feature time-sequence groups are created. The acoustic energy `RMS` is modeled by the average square root of the amplitude each ten millisecond, calculated with Octave (https://www.gnu.org/software/octave/). The word presence indicator `WRATE` is a binary feature. The feature’s activation function is set to 1 from the onset to the offset of the word. Content word presence indicator `CWRATE` is similar to `WRATE`, and the feature returns 1 only if the part-of-speech (POS) tag of the word is among nouns, verbs, adjectives and adverbs. Two semantic feature groups `PAR` (for paradigmatic) `SYN` (for syntagmatic) takes the latent values extracted from the corresponding embedding for the duration of the word utterance.

### G. Encoding Regressors

Voxels are considered as independent units as in other projects [4]. The BOLD temporal signal associated with an arbitrary voxel *j* is modeled as a linear combination of activities elicited by non-semantic and semantic features (3). The *f*_*i*_s are the previously presented feature temporal sequences. For each voxel, five classes of encoding models are trained: `RMS`, `RMS+WRATE`, RMS+WRATE+CWRATE, RMS+WRATE+CWRATE+PAR, RMS+WRATE+CWRATE+SYN. The order of the feature groups are consistent across different semantic modeling and the feature regressors are orthonormalized to remove potential confounds in later contrast analyses. Here hrf is the hemodynamic function used in Statistical Parametric Mapping software (https://www.fil.ion.ucl.ac.uk/spm/), provided by the Python library `nistats` (https://nistats.github.io/) [42]. This function mocks the BOLD response to neural activations. The convolution of hrf with feature activation functions models the hereby elicited BOLD signal, converts the activation function into *regressors* which are used in later regression/encoding. The coefficients *β*_*i,j*_ are to be determined via regression models.

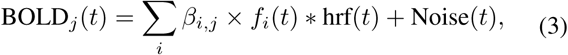

One particularity for `PAR` regressors is the filtering by regressor variance as they are based on PCA factored components. Regressors are dropped if its variance over the time-sequence is smaller than 10^−5^, which resulted 100 residual `PAR` regressors.

### H. fMRI Encoding

The relatively small observation samples collected from fMRI sessions and abundant regression features motivate the regularization with Ridge, which is described by (4) to promote the generalizability of the learned transformation matrix. The Ridge regression training algorithm uses a regularization coefficient *α*_*j*_ to balance the complexity of the model and the signal loss between the proposed model (the deterministic part as described by (2)) and the real signal BOLD_real,*j*_) measured for a given voxel *j*. For model classes containing `PAR` or `SYN` features, feed-forward feature selection is applied to maximally avoid potential overfitting by taking only the first *N*_*j*,cls_ number of features into consideration in voxel modeling. The values of the regularization coefficient *α*_*j*_ and *N*_*j*,cls_ are determined independently for each voxel via a grid search in a later stage of computation.

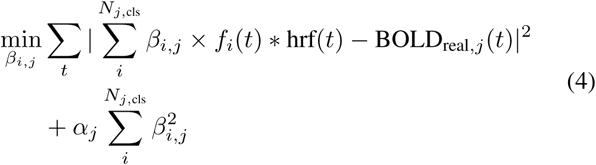

Models are trained and evaluated by means of a block-wise cross-validation where an fMRI record block is left to test the model. The coefficient of determination r2 is is a measure of the quality of the prediction. For each voxel *j* of a subject, the performance of the voxel model is averaged over nine cross-validations. Then, for each model class *cls*, the grid search retains only the combination of *α*_*j*_ and *N*_*j*,cls_ yielding the highest r2. Group scores are obtained by averaging the same voxel model scores of all subjects.

Voxel models are contrasted across model-classes and 3D maps summarizing voxel-wise differences reveal each feature’s spatial distribution in the human brain for language processing tasks. Five contrasts are calculated (see (5)) to show the evolution of lower-to higher-level language processing in the human brain. The first four contrasts are computed with nested voxel models. For the fifth, a non-nested model comparison framework [43] is experimented for regression simplicity. The contrast maps are visualized with the help of `nilearn`[42].

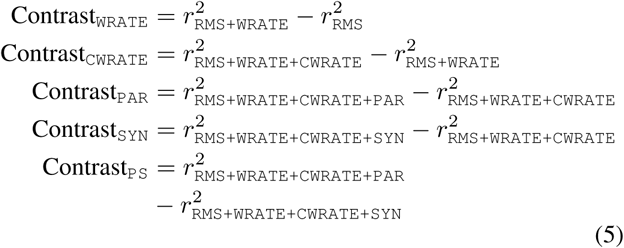

## III. RESULTS

### A. Twofold Dissociation in Embeddings

The constructed paradigmatic and syntagmatic embeddings successfully dissociates two semantic principles as shown by the word-pair semantic proximity ranking task (Table II), measured with paradigmatic-/syntagmatic-specific benchmarks. The French benchmark data are translated [44] from two widely used English datasets [45], [46] ^1^. By paradigmatic sub-graph curation, the French `PAR` contains only paradigmatic semantic information. The constructed French `SYN` significantly reduced paradigmatic information compared to the original embedding, while its syntagmatic counterpart is also slightly impacted, as indicated by the task results. To visually control the embedding quality, we sampled a few tokens from *the Little Prince* and examined the vectorial neighbours with TensorFlow (http://projector.tensorflow.org/) embedding projector from different embeddings and were reassured of the axis-wise purity of each embedding, thus the effect of axis dissociation. To further validate the methodology, the embedding construction and benchmarking are replicated in English using WordNet and a pre-trained GloVe embedding^2^. The replication achieves a better dissociation of two semantic axes, supporting our methodology, and suggesting that the French embeddings could be further tested with more appropriate behavioral benchmark data.

**TABLE II.**
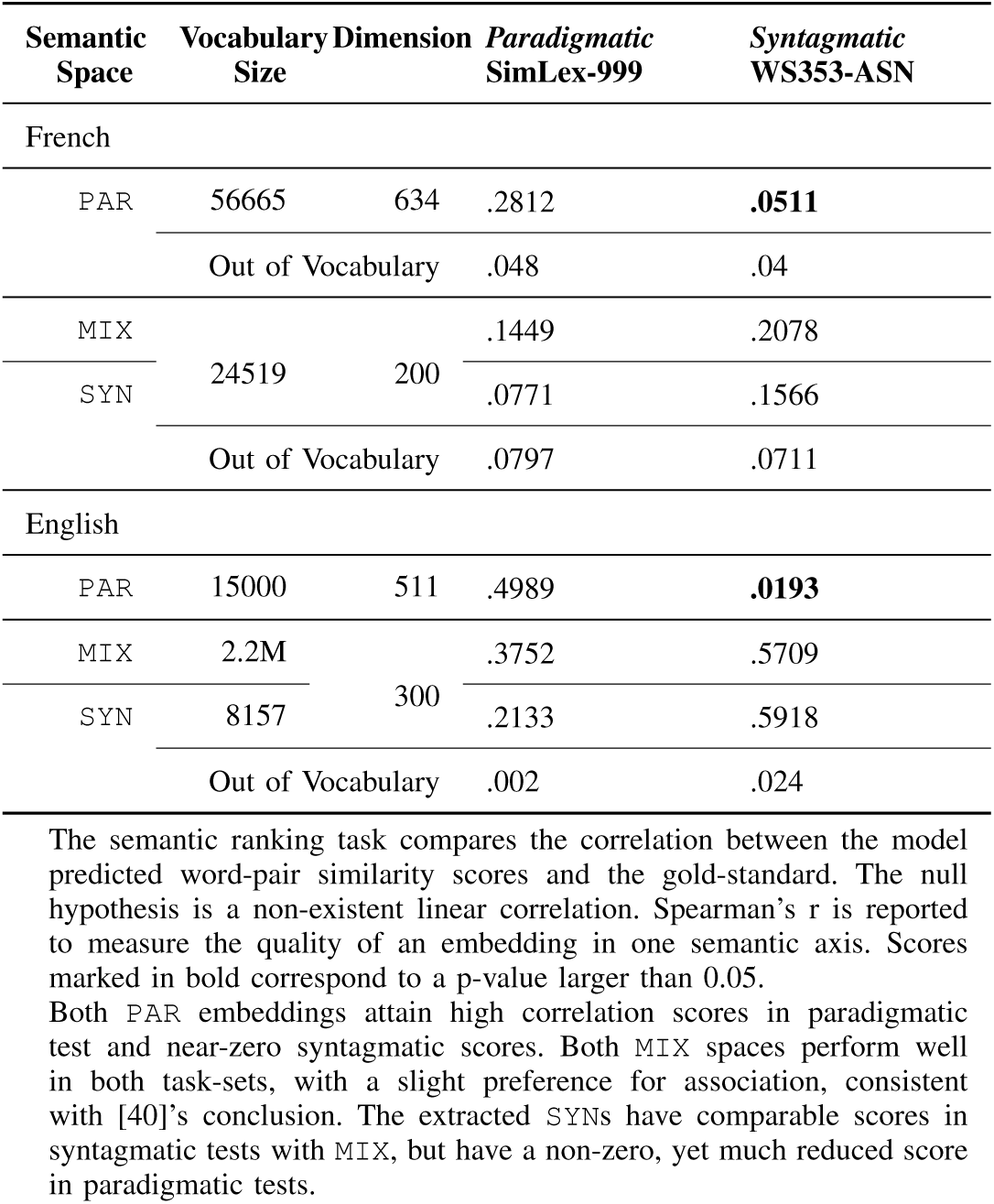
Semantic Space Semantic Ranking Task Results

### B. Primary Auditory Processing

With acoustic feature `RMS`, the r2 distribution map illustrated with obtained voxel model performances (Fig. 1) reveals several voxel-clusters with the most significant and largest activations from left posterior superior temporal gyrus (pSTG, Brodmann Area (BA) 41, peak at -60 -12 4, *r*2_*max*_ = 0.0362), followed by right pSTG (61 -13 2, *r*2_*max*_ = 0.0328). Other important yet much weaker activations are found in right middle cingulate cortex (MCC, BA23, 0 -24 29, *r*2_*max*_ = 0.0189) and right middle frontal gyrus (MFG, BA10, 29 56 20, *r*2_*max*_ = 0.0180).

**Fig. 1.**
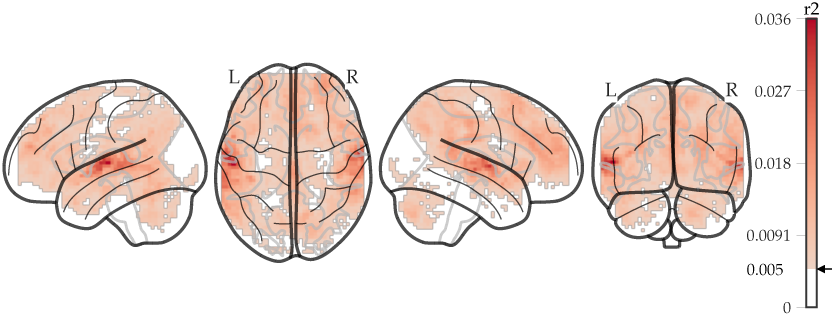
Acoustic energy feature reveals bilateral primary auditory cortices with a slight left lateralization.

### C. Gross Semantic Processing

Possibly due to the collinearity between `RMS` and `WRATE`, the addition of word presence indicator to the feature set does not improve voxel-models. The results are omitted. `CWRATE` on the contrary, boosts a large portion of voxels’ performance especially for those initially badly predicted with only `RMS` and `WRATE` features (Fig. 2), indicating that voxels processing semantic information has a limited usage of primary acoustic features.

**Fig. 2.**
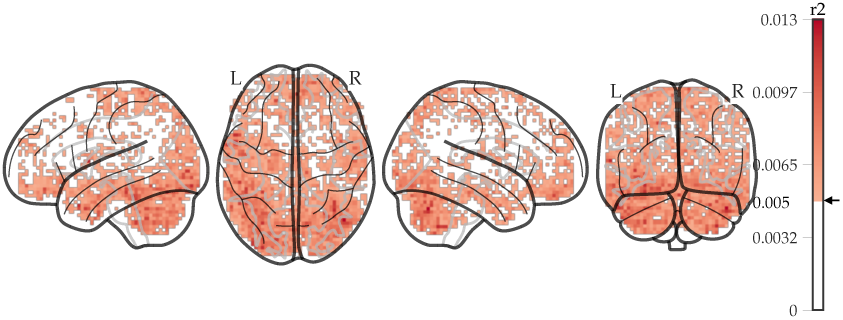
Content word presence indicator feature improvements are mainly located in bilateral temporal pole, inferior temporal gyrus, frontopolar prefrontal.

The major improvement of `CWRATE` is remarked in left middle and inferior temporal pole (TP) (BA38, -53 11 -33), bilateral posteroinferior temporal gyrus (ITG, including fusiform gyrus FG, BA19/37, -47 -43 -24, 45 -69 -38), frontopolar prefrontal cortex (fpPFC, near rectus gyrus, -5 46 -26) and posterior cerebellum (Wilcoxon’s W=136, Δr2>0.0067, p-value<10^−3.66^ uncorrected).

### D. Paradigmatic Processing

Fig. 3 illustrates the voxel performance improvements when paradigmatic semantic embedding features `PAR` are added to RMS+WRATE+CWRATE regressors. The most improved voxel clusters are located in bilateral middle temporal gyri (MTG, -51 -34 1, 57 -36 0, left cluster more significant), right angular gyrus (AG, 35 -65 44) and left superior parietal lobule (SPL, -27 -72 44) (W=210, Δr2>0.0079, p-value<10^−4.35^ uncorrected).

**Fig. 3.**
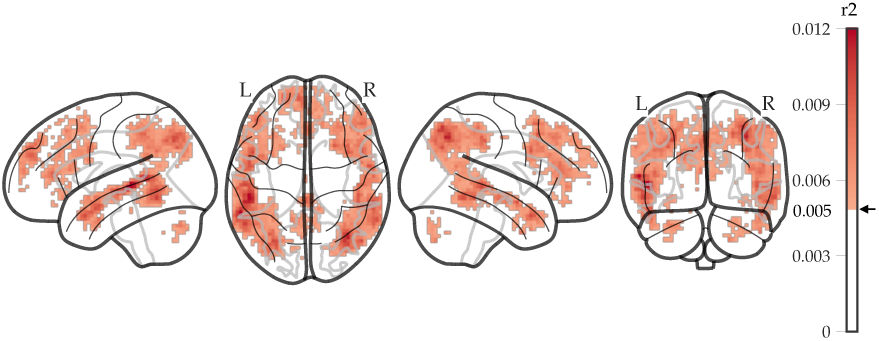
Paradigmatic embedding improves voxel models located in bilateral middle temporal gyri, angular gyri and left superior parietal lobule.

### E. Syntagmatic Processing

Fig. 4 illustrates the voxel performance improvements when syntagmatic semantic embedding features are added to RMS+WRATE+CWRATE regressors. The most significant improvements are slightly different from paradigmatic neural responses. The most significantly improved cluster, which is located in left MTG, is more posterior than that found for `PAR` contrast (−53 -58 3). Other important clusters include bilateral inferior frontal gyrus pars triangularis (IFGtri, -46 37 11, 50 35 8), left middle occipital gyrus (MOG, -32 -78 41), right AG (43 -75 39) (W=190, Δr2>0.0065, p-value<10^−4.18^ uncorrected).

**Fig. 4.**
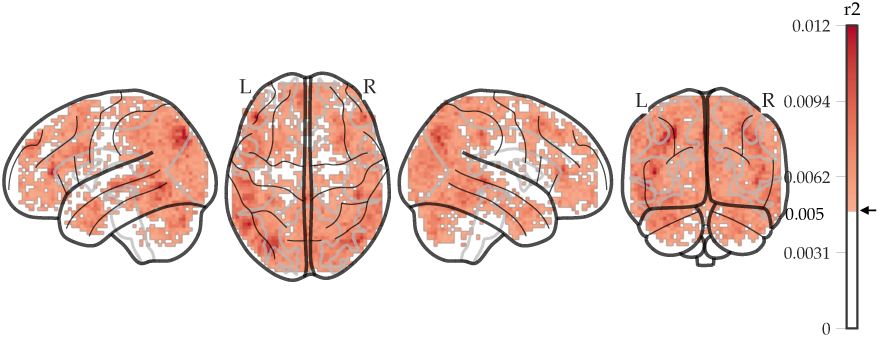
Syntagmatic embedding improves voxels from the bilateral anterior temporal lobe, posterior middle temporal gyri, bilateral inferior frontal gyri, right angular gyrus and left middle occipital gyrus.

### F. Paradigmatic and Syntagmatic Particularity

Current data suggest that paradigmatic and syntagmatic semantic processing are not supported by macroscopically dissociated neural components: the voxels along left MTG, right anterior and posterior MTG, bilateral SPL and IFGtri are improved equally by `PAR` and `SYN`.

To reveal a differential map for two semantic axes’ preference, the two classes of voxel models are contrasted with each other (Fig. 5). Most voxels are slightly better modeled by `SYN` models. Along the non-nested model comparison framework [43], the collinearity between paradigmatic and syntagmatic regressors are discovered. With `PAR` feature regressors, it is not possible to predict `SYN` features via a generalized linear model, whereas `SYN` regressors can be mapped to partially reconstruct first 5 `PAR` regressors.

**Fig. 5.**
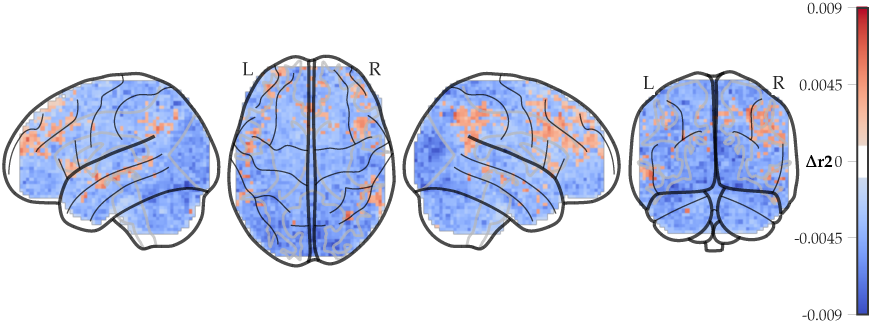
Paradigmatic and syntagmatic voxel model contrast. Positive values correspond to preference for paradigmatic processing, negative values to syntagmatic.

The Wilcoxon test on voxel models of 20 subjects yields two small significant voxel clusters for `PAR` in left superior frontal cortex (SFC, BA10, -27 59 25) and left anterior cingulum cortex (ACC, -8 34 25) (Δr2>0.0069, p-value<0.05 voxel-wise multi-comparison corrected). Left aSTG, right pSTG, right posterior superior temporal sulcus (pSTS, BA22) and left aSTS/middle TP are also reported (p<0.001 uncorrected). `SYN` found 17 small clusters in bilateral visual association areas (BA18), primary visual areas (BA17), ventrotemporal areas (ventral ITG, parahippocampal gyrus), left SPL, left thalamus and bilateral cerebellum (Δr2>0.0068, p-value<0.05 voxel-wise multi-comparison corrected).

## IV. DISCUSSION

The acoustic signal regressor is able to provide a primary auditory processing locus located in bilateral BA41, consistent with existing literature [47]. The slight left lateralization of acoustic voxel-model performance is consistent with Tervaniemi and Hugdahl’s [48] finding that left lateralization is linked to speech processing. The bilateral inferotemporal improvements, starting from fusiform gyri to anterior temporal regions, brought by content word presence indicator is associated with semantic memory processing [49], [50] and word-wise semantic priming effect in word meaning comprehension [51].

The spatial patterns of model improvements by paradigmatic and syntagmatic features in MTG interlace with each other, showing a gradient trend for both axes’ processing in MTG. Both features improve bilateral posterior MTG voxel models. This region is associated with word-meaning access across modalities and categories of concept [50] and was also proposed as a semantic hub [52]. IFGtri, improved by both semantic axes as well, seems to be associated with general semantic processing [53]. ITG, predominantly improved by syntagmatic embedding, is called by [54] as a *unification space* enabling syntactico-semantic integration. It suggests that syntagmatic axis is used in this process. Further investigation on the feasibility of syntactic prediction via only syntagmatic and the sentential semantic integration process is needed.

On paradigmatic/syntagmatic preferential contrast, despite the small effect sizes, the trends are roughly symmetrical across two hemispheres and the distribution is spatially regular. However, no region-of-interest (ROI) level analysis has revealed regional preference for paradigmatic processing, indicating no macroscopic neural structures related to the specific semantic axis on a group level.

Another paradigmatic embedding, namely the projection of paradigmatic embedding onto the mixed space has also been contrasted with existing `SYN` models. The contrasts shown by the two paradigmatic embeddings against `SYN` are globally consistent, but are not stable on the voxel level. The intersection of two paradigmatic-predominant regions (obtained by the `PAR`/`SYN` contrasts) is restrained to two small voxel clusters located in left posterior to middle STG/MTG and right inferior parietal lobule.

In conclusion, the paradigmatic/syntagmatic separation in semantic fMRI encoding leads to minute regional contrasts for both semantic axes. Syntagmatic processing is dominant in ITG and MOG. The ITG aspect could be related to semantic unification process. MOG’s linkage to imaginability[55] and its role in syntagmatic processing remain to be further investigated. The paradigmatic preference found in MTG is neighbored by syntagmatic counterpart, and in MTG both axes improve significantly model performance, indicating a gradient yet mixed nature of MTG semantic processing. The current results do not reveal a statistically established preference for the syntagmatic processing axis over the paradigmatic one and vice versa in the bilateral ATL, contrary to what might be expected following [18]. Future research could bridge syntactic processing, syntagmatic semantic processing and semantic composition. Distributional and structural neuroscience studies could also complete this current fMRI investigation by providing anatomical theoretical constructs.

## AUTHOR CONTRIBUTIONS

Songsheng Ying: Methodology, Software, Formal analysis, Data Curation, Writing - Original Draft, Visualization.

Sabine Ploux: Conceptualization, Methodology, Validation, Resources, Writing - Review & Editing, Supervision, Project administration.

## ACKNOWLEDGMENT

We thank Christophe Pallier and Snezana Todorovic, from the INSERM Cognitive Neuroimaging Unit (www.unicog.org), for making the fMRI data available to us, as well as some data analysis scripts. We also acknowledge the help of Laurent Bonnasse-Gahot, from CAMS-EHESS, for the conception of the semantic space dissociation algorithm and the fMRI analysis. Sincere thanks to Jialiang Lu, Jean-Pierre Nadal, Yang Yang, Xiaoqing Jin and Junyan Li for their efforts to make this research project possible.

## Footnotes

1 The translations are available on https://github.com/nicolasying/Similarity-Association-Benchmarks, commit `c97583f`.

2 The pre-trained model can be downloaded from https://nlp.stanford.edu/projects/glove.

